# Altered Lighting Conditions Elicit Sex-Specific Circadian Behaviors in Diurnal Grass Rats

**DOI:** 10.64898/2026.06.17.732698

**Authors:** Jeremiah Hartner, Noah Muscat, Mujtaba Khan, Katrina Linning-Duffy, Diksha Zutshi, Nicolette Ognjanovski, Lily Yan, Brendon O. Watson

**Affiliations:** Watson Laboratory, Department of Psychiatry, University of Michigan, Ann Arbor, MI, USA; Yan Laboratory, Department of Psychology, Michigan State University, East Lansing, MI, USA

**Keywords:** Circadian rhythms, Nile grass rat, Arvicanthis niloticus, diurnal, Seasonal Affective Disorder, Major Depressive Disorder, actigraphy

## Abstract

Circadian rhythms are crucial to biological functions, and cognitive functions such as attention, choice, and preference-related behaviors are modulated by circadian rhythms and disrupted in mood disorders such as Seasonal Affective Disorder (SAD) and Major Depressive Disorder (MDD). These neuropsychiatric diseases can be induced or worsened by alterations to daily light patterns and can also be treated with circadian-timed bright-light therapy, suggesting modulatory effects of light brightness on mood and behavior. While most laboratory rodents are nocturnal, the Nile grass rat (Arvicanthis niloticus) is diurnal, offering a unique model to study light modulation effects relevant to humans. In this work, we track daily activity in male and female grass rats under varied lighting for several weeks, revealing sex-specific circadian patterns and responses. These findings establish a foundation for mechanistic studies of light effects on mood-related brain circuits in diurnal animals.

## 1 Introduction

Circadian rhythms are crucial to basic daily functions, and disruptions or alterations to these rhythms are implicated in changes to brain health seen in neuropsychiatric disease states. Numerous functions, including cognition, attention, emotionality and preference-related behaviors are modulated by circadian rhythms (1–4). Further, Seasonal Affective Disorder (SAD) and Major Depressive Disorder (MDD) show a strong circadian rhythm in symptomology and can also be worsened or induced by disturbance in circadian cycles or light alterations (5–10). In addition, circadian-timed light therapy can be an effective treatment for both SAD (11,12) and non-seasonal depression, suggesting modulatory effects of light on mood regardless of season (9,12). This also suggests that brain circuits are modulated over the 24-hour cycle, which we have recently shown electrophysiologically (13), and are also affected by light, especially mood-related regions such as the medial prefrontal cortex (mPFC). Elucidating how mood-related circuits are impacted by circadian rhythms and daily light exposure is crucial to understanding human circadian brain biology, but methods to study those detailed dynamics are too invasive to use in humans, and so rodents provide a valuable experimental model.

Efforts to understand the biology of daily light rhythms in a manner most relevant to humans are hindered by the fact that commonly used laboratory rodents are nocturnal while humans are diurnal. While the central molecular circadian pacemaker is conserved in both function and phase between nocturnal and diurnal animals, in diurnal animals, light promotes wakefulness/arousal and can be therapeutic in humans (11, 12); while in nocturnal animals, light induces sleep (14). Though nocturnal animal behavior generally appears to be anti-phase to diurnal animal behavior in response to light exposure, identical stimuli can elicit different behavioral responses between diurnal and nocturnal animals (15). Further, the organization of circadian oscillator systems in nocturnal versus diurnal species differs in a more complex manner than a simple flip in the daily behaviors despite in-phase SCN pacemakers, as many but not all extra-SCN oscillators are phase reversed (16–18). Thus, studies of nocturnal models cannot be directly translated or simply reversed to understand functions of diurnal brains and their response to altered lighting conditions. In fact, numerous therapeutic agents proven effective in nocturnal rodent models failed in human clinical trials (19,20), which reveal a translational gap in our understanding of the neurophysiologic effects of light on diurnal brains.

Additionally, rodent research has prioritized the use of male-only data, generally due to the difficulty of controlling for the female estrous cycle, or pooled data of males and females. This approach fails to address the sex-specific differences seen in humans, where females are 1.6-2.0x more likely to be affected by SAD, MDD and anxiety disorders (21–24). Men and women also show a difference in antidepressant efficacy and treatment response for MDD and anxiety disorders (25–27). This is likely due to many factors, including but not limited to known differences in neurobiological factors like brain function (hippocampus, amygdala, and prefrontal cortex), hormonal influences, and immune and stress response systems (28–31). These observations indicate that independent scientific exploration of the biology of both sexes may be key for translation to human disease.

The Nile grass rat (NGR; *Arvicanthis niloticus*) is diurnal in the wild (32) and is now a well-established laboratory model exhibiting diurnal patterns of sleep, locomotor activity, mating behavior, body temperature and secretion of hormones in the laboratory (33–36). Further, NGRs show reduced daytime behavior in lower light and have shown characteristics consistent with SAD in reduced lighting conditions (37,38). However, sex-specific 24-hour behavior patterns in various lighting conditions have not been examined. This is especially important given the contribution of circadian rhythms and light to mental health (1–4), and the known human sex differences in susceptibility to anxiety disorders, MDD, and SAD (22,39–42). SAD and MDD are both responsive to bright light therapy but require more mechanistic study in a diurnal animal model. Like patients, NGRs also show brightness-related behavioral alterations (43) and in fact standard laboratory lighting conditions, which provide much less light than natural sunlight may induce depressed behavior. For example, NGRs limited to a dimly lit environment in the daytime exhibit reduced sweet solution preference behavior (44,45), while NGRs exposed to bright light in the daytime show improved spatial memory (46,47), indicating that NGR physiology and behavior are similarly affected by variations in lighting conditions. However, there have not been attempts to quantify at both female and male behavior, nor have there been characterizations of movement over longer time scales involving continuous recordings over many months in varying lighting conditions.

In this project, we quantify the daily actigraphy of both male and female diurnal rodents over extended periods of time in 1) standard laboratory lighting conditions (300 lux in daytime) 2) more naturalistic/brighter light (1000 lux daytime) and 3) constant darkness (<10 lux) “free-run” conditions to assess daily and circadian rhythmicity. We quantify results by tracking changes in videographic pixel values from frame to frame. We aim to reveal the rhythms of daily activity across lighting conditions in diurnal mammals using a novel tracking technique to improve our understanding of behavior in rodents that share diurnality with humans. This can in turn establish a novel experimental paradigm to study the biology of SAD and/or light therapy.

## 2 Materials and Methods

### 2.1 Subjects

Male and female Nile grass rats (Arvicanthis niloticus) were acquired from the well-established breeding colony at Michigan State University managed by Dr. Yan (30). Animals were transported to the University of Michigan at 8-12 weeks of age and allowed to acclimate to the new facility for a minimum of 3 weeks prior to testing. Animals are initially group-housed in standard Plexiglas cages (17” x 9” x 8”) and provided nesting materials and hay for enrichment. Animals are transferred to single housing a minimum of one week prior to experiment start.

Animals were housed with a 12:12hr light-dark cycle under standard vivarium lighting (300 lux as measured from the center of the cage on the top-rack of vivarium shelving <4 feet from the overhead ceiling lights) with water and food (Prolab RMH 2000 5P06, Brentwood, MO, USA) ad libitum. Our vivarium lighting measures 100-300 lux depending on location of the animal cage within the housing rooms relative to the overhead lighting. The animals in this study were all housed on upper racks with a light intensity of 300 lux, which is consistent with an assessment of other research institution typical vivarium lighting conditions (Dauchy, Patel). For our bright light conditions, we added LED bulbs above the animal racks to raise the light intensity to >1000 lux with the goal of having the animal experience light intensities more like what a wild NGR might experience in outdoor lighting conditions considering direct sunlight is generally obscured by NGR tendency to burrow in tall grasses or soil.

Male and female NGRs were randomly assigned a cage position on the top rack of the vivarium shelving, and standard cage tops were modified using wire mesh (304 stainless steel, plain weave, 1.3mm aperture, 0.15mm wire diameter) to allow overhead camera capture with clear visibility of the animal. Four animals in four individual cages were recorded continuously at 12 frames per second 24 hours per day using StreamPix software (Norpix, Montreal, CA) paired with monochrome cameras (Ace U acA1300-200um, Basler, Exton, PA, USA) with 6mm, ½” lenses (Edmund Industrial Optics, Barrington, NJ, USA) and bandpass near infrared filters: 845 to 860 nm (Midwest Optical Bi850-43, Graftek Imaging, Austin, TX, USA). for the duration of the experiment. Camera was triggered to record for 59:59 minutes at a time and then re-started every 60 minutes.

Each cohort of animals was recorded over a period lasting 2-3 months, beginning with 4 weeks in 300 lux standard vivarium lighting conditions during lights-on phase of a 12:12-hr light-dark cycle, followed by 4 weeks of exposure to 1000-1300 lux LED lighting (CNSunway Lighting, Shenzhen, CN) only during the lights-on period of the 12:12-hr light-dark cycle, and finishing with 4 weeks in winter-like dim lighting at 0-10 lux (Full Dark; FD) achieved by enclosing the cages under blackout curtains for 24 hours per day to mostly shield the animals from exposure to the vivarium room lighting operating on a 12:12hr light-dark cycle at ∼300 lux during the lights-on phase (Fig. 1). The experiment consisted of 3 cohorts of 4 animals each for a total of 12 animals (6M/6F). The first cohort (2M/2F) were not exposed to the FD condition. The 2nd and 3rd cohort were exposed to all lighting conditions (Table 1). All recorded experimental days from all animals were fully analyzed and included in the results. All procedures were conducted following the guidelines set by the Institutional Animal Care and Use Committee (IACUC) of the University of Michigan.

**Figure 1:**
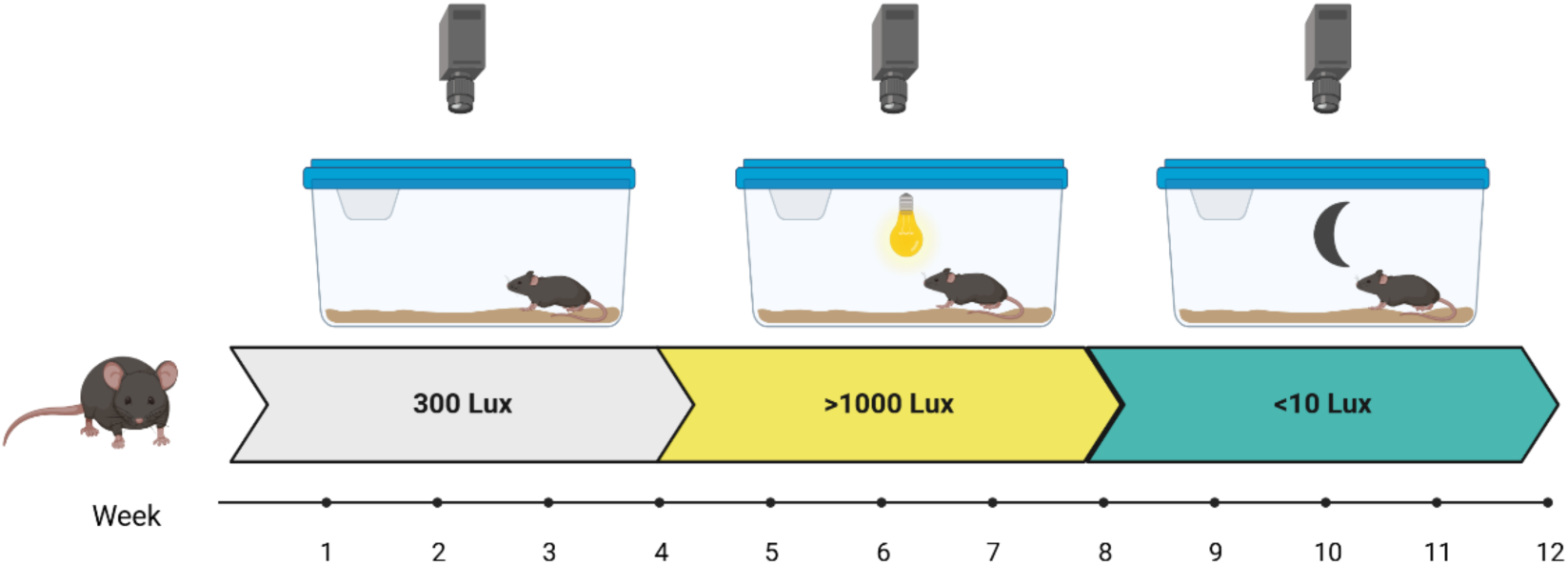
Experimental timeline. Schematic of recording protocol over the course of 3 months. Some animals completed all 3 phases of the protocol (n=8), a subset were recorded in only 300 lux and 1000 lux conditions (n=4). Created in BioRender. Hartner, J. (2026) https://BioRender.com/ybmrx7e

**Table 1:** Experimental Design by Animal.

| <u>Animal ID</u> | <u>Sex</u> | <u>Cohort</u> | <u>Rack Location</u> | <u>Condition 1:</u><br><u>300 lux</u> | <u>Condition 2:</u><br><u>>1000 lux</u> | <u>Condition 3:</u><br><u><10 lux</u> |
| --- | --- | --- | --- | --- | --- | --- |
| AO1 | M | 1 | Left | Y | Y | N |
| AO2 | M | 1 | Mid Left | Y | Y | N |
| AO3 | M | 1 | Mid Right | Y | Y | N |
| AO4 | F | 1 | Right | Y | Y | N |
| AO5 | F | 2 | Left | Y | Y | Y |
| AO6 | F | 2 | Mid Left | Y | Y | Y |
| AO7 | M | 2 | Mid Right | Y | Y | Y |
| AO8 | F | 2 | Right | Y | Y | Y |
| AO9 | M | 3 | Left | Y | Y | Y |
| AO10 | F | 3 | Mid Left | Y | Y | Y |
| AO11 | F | 3 | Mid Right | Y | Y | Y |
| AO12 | M | 3 | Right | Y | Y | Y |

### 2.2 Actigraphy

Actigraphy analysis was performed by developing Python-based code (https://github.com/BrendonWatsonLab/ActigraphyWork/tree/main/ActigraphyCode) to extract number of pixels showing changes in brightness over time, frame-to-frame, and was used as a proxy for animal movement. To achieve this, a simple script was created with several steps including grayscale conversion, dual-threshold filtering (global thresholding and percentage change thresholding), dilation, and region size filtering. This script tracks only pixels that exceed the thresholds for each step (details below), requiring that spatially contiguous groups of a minimum number of pixels show change in brightness, which minimizes noise and selects changes likely due to animal movement. The output provides a Selected Pixel Difference (SPD) value, which is the count of all pixels per frame that meet the selection criteria. By incorporating timestamps with each video frame, we then use SPD as a proxy for animal movement over time.

To compute the pixel-wise changes frame-to-frame, we first convert each pixel value from color to grayscale, which gives a single brightness value that can be more simply tracked frame-to-frame. We then perform dual-threshold filtering, where the change in brightness value must exceed both a global threshold and percentage change threshold compared to the previous frame. All pixels that meet both criteria are converted to binary (white or black), and only white pixels are kept. We then dilate areas of white pixels to fill in small gaps between neighboring white pixels. This creates multiple blobs - one for each area of the pixel intensity change - from which we can find the one corresponding to the moving animal. The size of each of the contiguous areas of white pixels is calculated and a user-defined threshold is used to eliminate smaller white regions which are unlikely to be representative of pixel changes due to animal movement. The final output is the SPD value representing the total number of white pixels in each frame.

Each of the thresholding parameters is user-defined through a graphical user interface (GUI), which allows for iterative testing to refine pixel change tracking. This is best achieved by using a second script to provide video output highlighting all selected pixels in each frame based on the user-defined parameters (Fig. 2A-B). Thus, the user can alter the thresholds for pixel changes and immediately assess the quality of animal tracking. Once pixel change threshold parameters are optimized, the script can be run on all videos within that same environment without further calibration, and further refinements can be made for different video environments or varying light conditions. Standard thresholds are…

**Figure 2:**
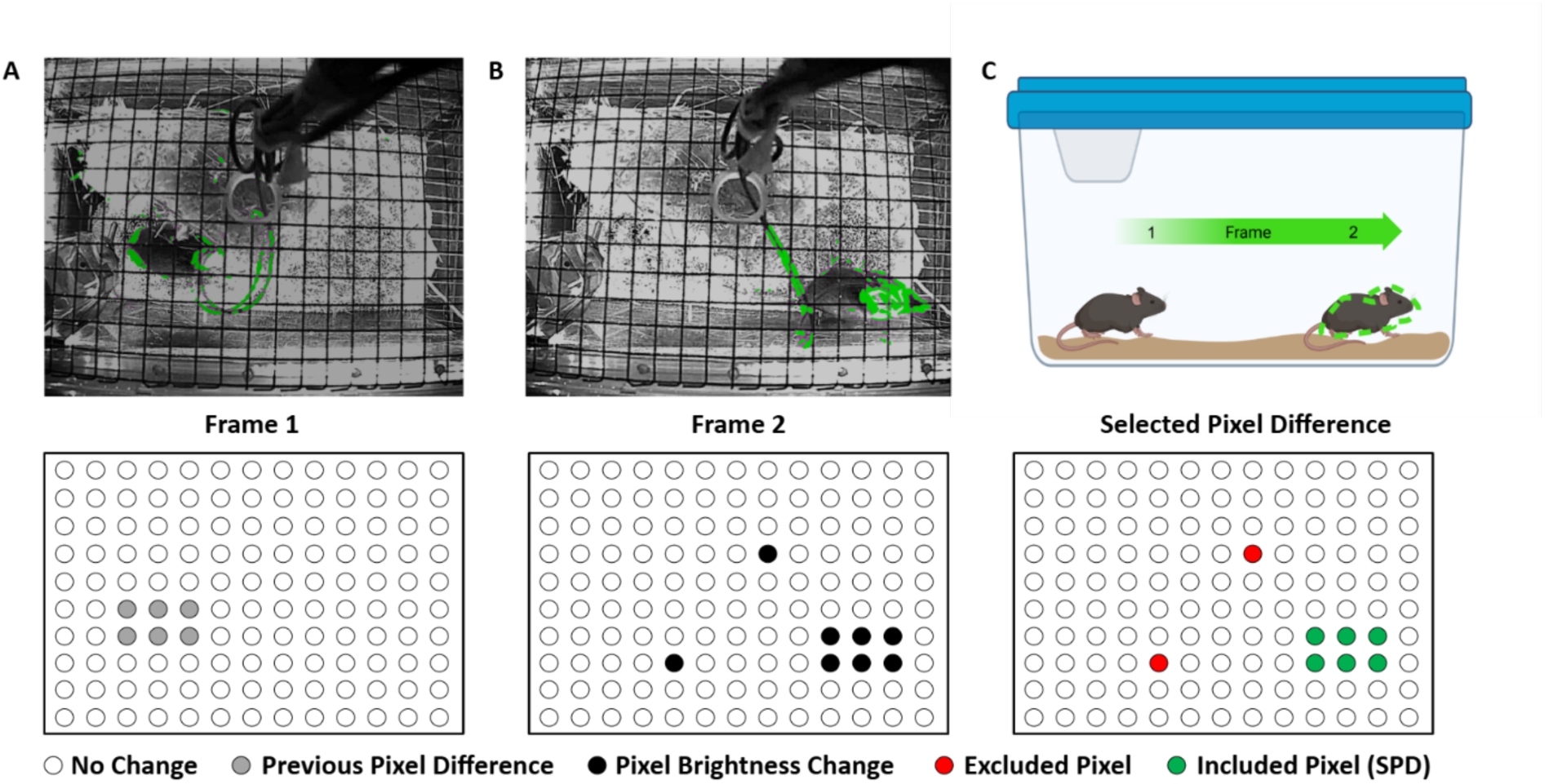
Selected Pixel Difference as a measure of animal movement. A) Top: Demonstrative example. Frame 1 shows the animal’s current position as tracked using frame-to-frame actigraphy. Frame-to-frame selected pixel changes are represented in green. Bottom: Schematic illustrating pixel changes within the captured camera frame. B) Top: Example frame 2 showing animal movement to the opposite side of the cage. Actigraphy SPD tracking represented in green. Bottom: Schematic illustrating pixel changes within the captured camera frame. C) Top: Representative output of SPD as the animal traverses the cage. Bottom: Schematic demonstrating exclusion of pixel brightness changes to refine SPD tracking of animal-related movement frame-to-frame. Created in BioRender. Hartner, J. (2026) https://BioRender.com/ybmrx7e

After calibrating parameters for multiple animals in different lighting conditions, a single set of parameters (pixel threshold = 30, area threshold = 100, dilation = 5) optimally tracked our animals’ movement over time irrespective of animal, cage conditions, lighting, or cage placement within the room (Fig. 2). Tracking of other animal types in varying environments may need different parameters, though this was not tested. These parameters were then applied to all videos from all animals in the experiment, allowing us to efficiently pull selected pixel difference from each frame of continuous months-long video recordings. We then used these pixel changes as a proxy for animal movement to compare hourly rhythms of activity across animals and lighting conditions over time. The output of the script is a single .csv file showing the SPD value for every time-stamped video frame for every animal across all conditions.

Data are output as .csv files and to enable locking to clock time and therefore subsequently lights-on Zeitgeber time, each frame is timestamped in POSIX timestamps based on video file start time plus frames elapsed multiplied by seconds-per-frame (i.e. 0.33s for 30 frames per second).

### 2.3 Statistical Analyses

Data analysis was performed using MATLAB (R2023a, MathWorks Inc.) to compare absolute SPD and normalized (z-scored) SPD values. All timestamps were converted to Zeitgeber time based on lights-on time for each animal. Changes in SPD were compared to a baseline/control condition to track changes in animal movement across minutes or hours for each of the lighting conditions. All diurnality analyses were performed by normalizing each animal’s hourly movement by averaging each hour of the day across the final 14 days of baseline 300 lux conditions (all values divided by this). For assessing changes to movement in varying lighting conditions, the SPD was normalized to the final 14 days of the bright lighting condition (1000 lux) and to the final 7 days of the FD condition (due to shifting of animals’ subjective day/night without light entrainment) and compared to the individual animal’s baseline movement in 300 lux. The final 14 days were chosen to allow the animal multiple weeks to acclimatize to the new lighting condition and control for any stress response likely to be seen immediately following an environmental alteration. The resulting datasets were examined by comparing binned averages between either raw SPD or normalized SPD in bins of 5mins or 60mins across days/weeks to determine significant differences in movement across time in different lighting conditions. Comparisons were made using a variety of t-tests and ANOVAs.

#### 2.3.1 Diurnality Index (DI)

A diurnality index was calculated to simplify activity differences seen during the lights ON phase compared to the lights OFF phase. DI is calculated by using the percentage change in raw (not normalized) SPD during lights ON compared to OFF for each of the lighting conditions:

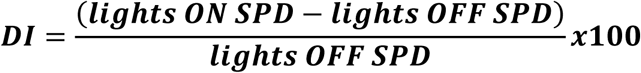

#### 2.3.2 Linear Mixed Effects (LME) model

To compare activity levels between lighting conditions while accounting for the repeated measures structure of the data, we used linear mixed effects models (LME) in MATLAB (fitlme) as an alternative to ANOVA due to the repeated measures of animals over varying numbers of days. Each model included lighting condition as a fixed effect and individual animal as a random intercept, allowing each animal to have its own baseline activity level. This approach uses all daily observations (approximately 14–30 days per animal per condition) while accounting for the non-independence of days recorded within the same animal. Model fit was assessed using Akaike Information Criterion (AIC). Fixed effects were tested using t-statistics with Satterthwaite degrees of freedom approximation as implemented by MATLAB’s fitlme function. For between-sex comparisons, sex was included as the fixed effect with individual animal as a random intercept.

## 3 Results

### 3.1 Daily Movement Patterns in Standard Vivarium Lighting

We first confirmed NGRs show diurnal patterns of movement in our laboratory setting over time. To test this, animals were moved to single housing for at least one week prior to testing to allow the animals to acclimate to that condition. A 12-week paradigm is used as shown in Figure 1 exposing animals to 300 lux, 1000 lux and <10 lux daytime light for 4 weeks each. Quantification of daily movement was done using the SPD metric outlined in the methods. A total of 12 single-housed NGRs (6 male, 6 female) were recorded in 3 separate cohorts of 4 animals each (Table 1).

To assess circadian rhythmicity, a Lomb-Scargle Periodogram was performed (plomb’ function in MATLAB) using 5 minutes bins of SPD for every 0.5 hours between 0.5 and 36 hours in the final 14 days of the baseline 300 lux condition. The last 14 days were chosen as the analysis window to account for any stress while acclimating to the new environment along with a noticeable settling of animal activity during the first 2 weeks (Fig. S3A). Individual animal activity was z-scored prior to pooling, and the resulting periodogram showed NGR activity was consistently cyclical with a large peak near both 12 and 24 hours, indicating a strong daily cycle of movement patterns (Fig.3A). Pooled actogram analyses of SPD for the last 14 days of standard laboratory lighting conditions of 300 lux (n=12 NGRs) show clear increases in activity during lights ON compared to OFF, with clear bands of power for SPD near ZT6 and ZT12 (paired t-tests, p < 0.05 for hours 5, 6, 10-12, 15-21, 23; Fig.3B). Average daily activity profiles were created using hourly averages of mean SPD for each animal over all five weeks to demonstrate sustained increases in SPD during lights ON (ZT 0-11) compared to lights off (ZT 12-23), indicating more hourly movement during the lights ON period with distinct peaks in activity around ZT 6 and ZT 11-12 (Fig.3C). Quantification of SPD also indicates significantly increased activity during the full lights ON phase (ZT 0-11) vs full lights OFF phase (ZT 12-23) in the final two weeks of the baseline recording (paired t-test, p=0.038; Fig.3D). Due to significant increases in SPD during the transition periods surrounding the switch from lights ON of OFF (ZT 11-12) compared to the full day (two-tailed paired t-test, p=0.0001; Fig.3D) and from lights OFF to ON (ZT 23-0) compared to full night (two-tailed paired t-test, p=0.008), as a more accurate representation of diurnality (as opposed to crepuscularity), we also compared the middle of the lights ON phase (ZT 3-9) to the middle of lights OFF during the final two weeks in 300 lux and saw significant increases in mid-day activity compared to mid-night (ZT 15-21; two-tailed paired t-test, p=0.001; Fig.3D). Weekly averages across 5 weeks of baseline recordings showed significant differences in both full day vs full night as well as mid-day vs mid-night animal activity for each of the 5 weeks (Fig. S1). These results indicate that NGRs display strong diurnal cycles of activity across many consecutive days of recording, in our laboratory vivarium environment.

**Figure 3:**
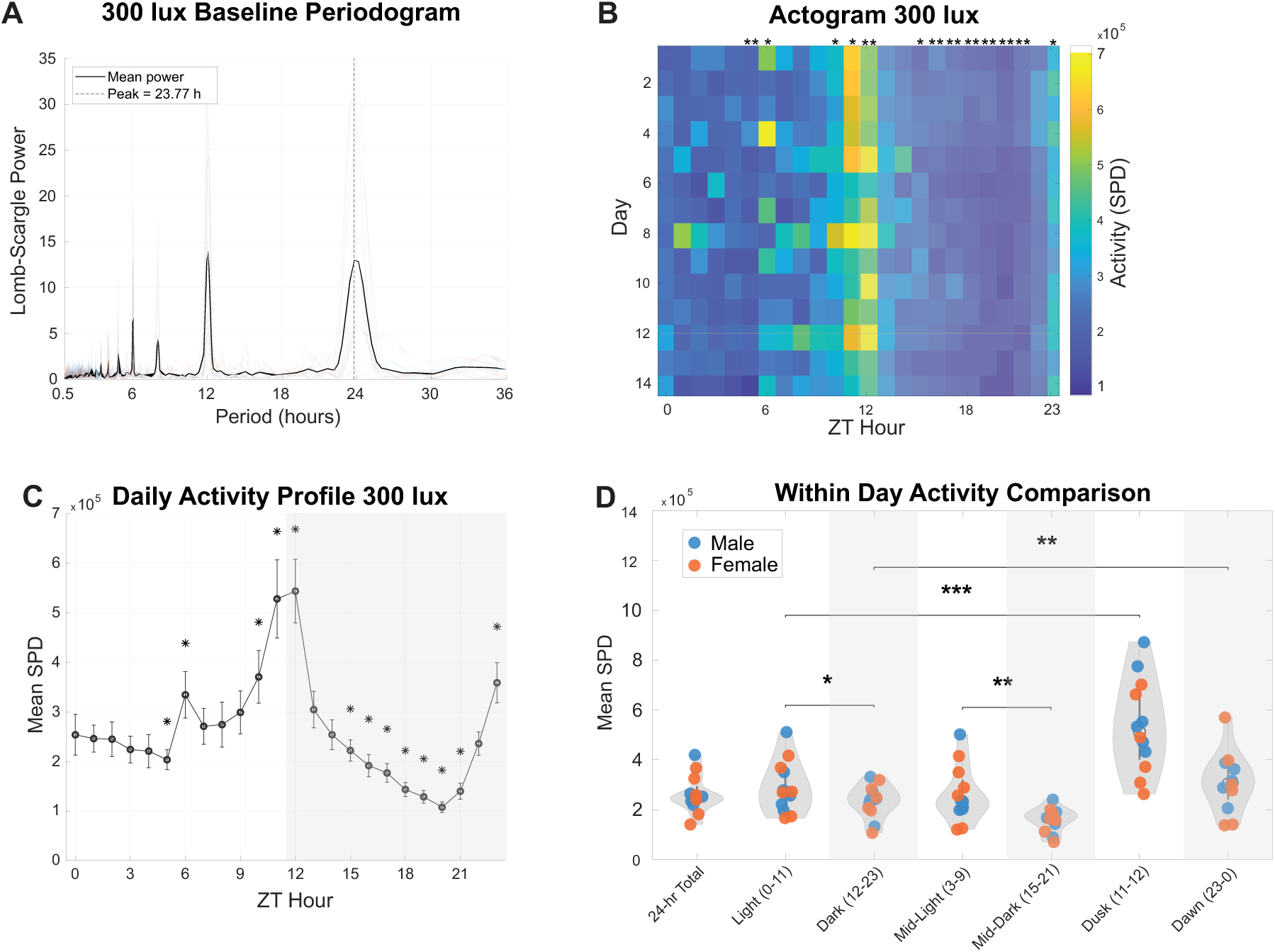
Characterization of animal movement using selected pixel difference (SPD). NGRs display diurnal patterns of activity. A) Periodogram testing for rhythmicity and patterns of movement across the final 14 days of baseline 300 lux conditions. Peaks indicate a repeating pattern of activity at a given lag. Animal activity shows a clear peak around 24hrs indicating patterned animal movement on a circadian scale. B) Actogram analysis detailing average pixel change by hour (n=12) across the final14 days of recording at 300 lux daytime light. 300 lux is considered standard laboratory conditions. Degree of pixel change representing animal activity demonstrated by heat-map (yellow indicates more pixel change). Left portion of plot (lights-on hours) generally shows more activity than right (lights-off hours). Asterisks above hourly columns indicate significant differences between that hour and the average of the other 23 hours; *= p < 0.05, ** = p < Bonferroni corrected threshold due to multiple t- test comparisons. C) Mean Selected Pixel Difference for all animals by hour across the final 14 days of the 300 lux baseline condition. Activity shows peaks in mean normalized activity at ZT6 and 12 as well as more activity during lights ON vs OFF (left vs right side), matching the actogram analysis. Asterisks indicate hours that differ from the distribution of all 23 other hours: activity peaks around ZT12 and reaches minimum around ZT20. D) Quantification of per-animal mean normalized SPD for different portions of the 24-hour cycle. Each dot is one animal, blue circles = male animals; orange = female. Numbers in parentheses indicate hours used for each period. Statistically significant differences are represented by asterisks.

### 3.2 Effect of Light Alteration on Daily Movement Patterns

Given the influence of light on mood in humans who are diurnal and since NGRs display diurnal activity patterns, we next tested the effects of increased daytime lighting on NGR actigraphy. Environmental lighting is commonly categorized into indoor versus outdoors based on a 1000 lux light intensity threshold (48–50), where the intensity of light below 1000 lux is generally experienced indoors. Following the baseline period, all 12 animals were exposed to 1000-1300 lux light during the lights ON phase for an additional 4 weeks. Bright lighting was achieved using overhead rack-mounted LED lighting with brightness measured at NGR eye level. A portion of the animals (Cohorts 2-3; n=8) were also then exposed to 4 weeks of full dark (FD, <10 lux) lighting after the 1000 lux condition (Figure 1).

To assess for behavioral activity pattern changes in response to light across the course of the day, we began by plotting mean hourly activity for all animals (n=12 male and female pooled) in the 300 and 1000 lux light conditions, using the final 14 days in each condition. We compared activity levels hour-by-hour across lighting conditions and found that NGRs show significantly more activity in the 1000-ON condition for each daylight hour except for ZT 11 (two-tailed independent t-tests, p<0.05 for 11 of 12 hours, Fig. 4A). The bright light has a smaller effect on animal activity during the OFF period where hourly average comparisons were significantly decreased compared to the 300 lux-OFF period during only 2 of the 12 hours, both of which fell near light transitions (p<0.05 for ZT 13 and ZT 23; Fig. 4A). While these effects are lower in amplitude and only occur for two hours, it is noteworthy that they are downward in direction rather than upward with increased lighting – such that increased daytime activity coincides with reduced nighttime activity. The effect on 1000-ON activity was quantified further by comparing 12-hr averages of lights ON and OFF for each condition, and bright light exposure significantly increased daytime NGR activity compared to daytime activity in the 300 lux-ON (two-tailed paired t-test, p=0.02; Fig. 4D), though the crepuscular-like increase in activity just prior to the lights OFF transition in 300 lux mitigates some of this effect. In line with the daily activity profile, the bright light did not significantly affect overall animal activity during the lights OFF phase (1000-OFF 12-hr average vs 300 lux-OFF 12-hr average, two-tailed paired t-test, p=0.58; Fig. 4D). Animals in the bright light condition not only maintained their diurnality (two-tailed paired t-tests, 1000-ON 12-hr average compared to 1000-OFF 12-hr average, p=0.002; Fig. 4D), but showed significant increases in diurnality index (DI) compared to 300 lux (paired t-test, DI 1000 vs 300 lux, p=0.012; Fig. 4F) – potentially via increased daytime activity and small tendency towards reduced nighttime activity.

**Figure 4:**
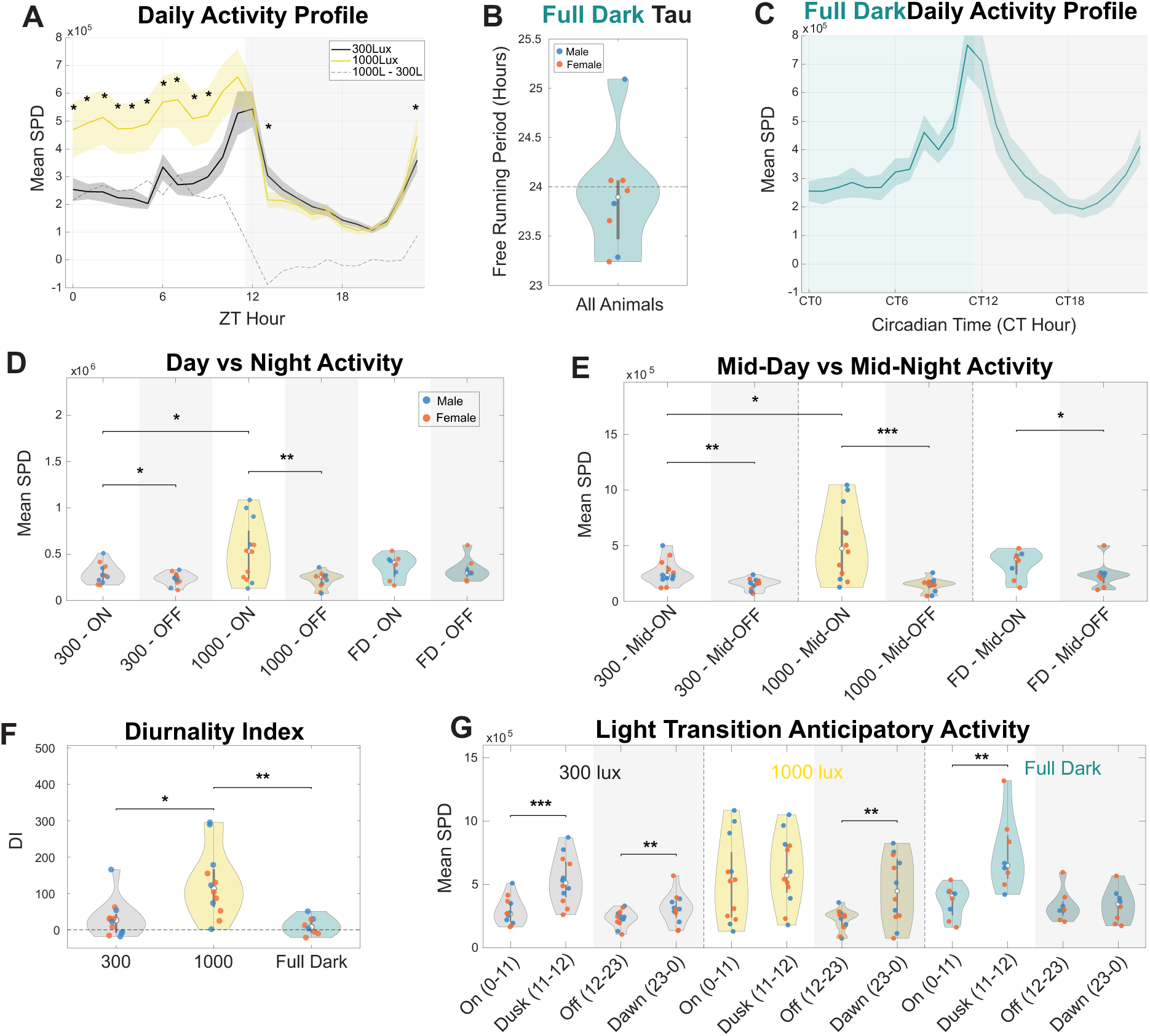
Bright light increases daytime activity and increases diurnality. A) Line plots of hourly averages of normalized SPD for 300 lux and 1000 lux conditions. Dotted line indicates change from 300 lux to 1000 lux (via subtraction of hourly means). Asterisks indicate significant difference between hourly means for 300 lux versus 1000 lux conditions. The increases in brighter light are concentrated in the light hours, but dusk and dark behavior are not significantly changed. B) Free running period length (Tau) for the last 7 days of all animals that were tested in full dark (n=8) as computed by a Lomb-Scargle Fourier transform. Dashed line indicates the expected 24hr rhythm. Male and female individual free running periods represented by blue and orange dots respectively. C) Full dark activity profile for all animals (n=8) in the last 7 days of the full dark condition. Each individual animal’s hourly means were re-aligned to CT time prior to averaging across animals. Pooled hourly means are represented by teal line with teal shading (SEM). D) Quantification of 12-hr lights ON and OFF averages for SPD by light condition (ZT 0-11 lights ON and ZT 12-23 lights OFF for the 300 lux and 1000 lux conditions, CT 0-11 for subjective day in full dark and CT 12-23 for subjective night in full dark). Male (blue) and female (orange) data is pooled per violin, and statistically significant differences between groups are represented by asterisks. Differences between day and night in both 300 and 1000 lux, but difference with 1000 lux is greater. E) Same as in D except analysis window is confined to the middle six hours of each phase (ZT 3-9 for lights ON, ZT 15-21 for lights OFF; full dark CT 3-9 for subjective day and CT 15-21 for subjective night). These results are qualitatively similar to full ON/OFF period analysis except FD condition now also shows difference between day and night. F) Diurnality Index (DI = (lights ON SPD – lights OFF SPD)/lights OFF SPD) x100) comparison for pooled sexes by light condition. G) Quantification of anticipatory peaks of activity during light-transition times (last hour before transition and first hour after transition, ZT/CT 11-12 and ZT/CT 23-0) compared to averages of 12-hour lights ON or OFF times (ZT/CT 0-11 and ZT/CT (12–23). Significant differences demonstrate the loss of anticipatory increase for Bright-Dusk and FD-Dawn, since earlier light phase values are higher.

To reduce crepuscular activity contributions in our quantifications, we tested mid-day vs mid-night and still saw a significant increase in daytime activity compared to nighttime in the 1000 lux condition (paired t-test mid-1000-ON vs mid-1000-OFF: p = 0.0008, N=12; Fig. 4E). We also examined the crepuscular time windows by quantifying differences in anticipatory activity during the final hour of lights ON through the first hour of lights OFF (ZT 11-12) with the full lights ON average (ZT 0-11) for each lighting condition. We found that in 300 lux there is a significant increase in activity just prior to lights OFF (paired t-test, 300-ON vs Dusk: p=0.0001 (N=12); Fig.4G), whereas in the bright light the anticipatory movement prior to lights OFF is not statistically different from the full lights ON period (1000-ON vs Dusk: p=0.356 (N=12); Fig.4G). This emerging lack of difference is likely due the large increase in activity during the mid-ON period with 1000 lux light while the crepuscular activity values are unchanged between 300 and 1000 lux (paired t-test, 300-Dusk vs 1000-Dusk: p=0.392 (N=12); Fig.4G). These findings indicate that brighter daytime lighting increases diurnality in NGRs due to increased activity during the lights ON phase.

Exposing the animals to the FD condition allowed us to observe their circadian pattern and calculate each animal’s free running period during the fourth week of FD using a Lomb-Scargle periodogram (Table 2). The average free running period is 23.90 hrs, not significantly different from 24hrs (one-sample t-test, n=8, p = 0.64; Fig.4B). However, because of the individual differences in free running period, some shorter some longer than 24hrs, we used only the final 7 days of this lighting condition for quantification because using longer periods would incorporate larger shifts in subjective time of day. To account for individual circadian phase difference following several weeks of free running in FD, we aligned each animal’s daily activity profile by circadian time (CT). To do this, we computed each animal’s daily activity profile by averaging hourly activity across the last 7 days of the FD condition, applying a 3-hour circular moving average, and identifying the peak activity hour. We found that for each of the 8 animals in the full dark environment, the daily activity profile displayed two prominent peaks of activity, directly mirroring the animals entrained by light, only shifted in time (Fig. S2). Because light-entrained animals in our study all showed the most prominent peak of activity at ZT11 (Fig. 4A), we re-aligned each FD animal’s daily activity profile by applying an integer-hour phase offset to shift each animal’s most prominent peak to CT11 (Fig. S2) prior to pooling and averaging the data for quantification (Fig. 4C). This re-alignment to circadian time allows us to quantify each animal’s activity during the subjective day/night despite the individual variations in perceived time of day.

**Table 2:** Free-Running Period (tau) During Fourth Week of Full Dark by Animal.

| <u>Animal ID</u> | <u>Sex</u> | <u>Cohort</u> | <u>Tau</u> |
| --- | --- | --- | --- |
| AO5 | F | 2 | 23.24 |
| AO6 | F | 2 | 24.07 |
| AO7 | M | 2 | 23.29 |
| AO8 | F | 2 | 23.66 |
| AO9 | M | 3 | 25.09 |
| AO10 | F | 3 | 23.96 |
| AO11 | F | 3 | 24.07 |
| AO12 | M | 3 | 23.83 |

Interestingly, animals in the fourth week of full dark still showed significantly more activity during the middle of the subjective lights ON period compared to the middle of the subjective lights OFF period (paired t-test, ZT3-9 vs ZT15-21, p=0.013; Fig. 4E) despite the lack of light entrainment. Further, all FD animals still showed anticipatory increases in activity (paired t-test, FD-ON vs Dusk, p=0.003; Fig. 4C/G), indicating that these animals naturally follow a diurnal pattern of activity with expectation of light transition. Due to re-alignment of daily activity based on the larger peak at CT11, the anticipatory peak amplitude obscured the 12-hr comparison of subjective day vs night (paired t-test, FD-ON vs FD-OFF, p=0.224; Fig. 4D). These animals also showed a small decrease in diurnality index as compared to the 300 lux conditions, though not significant (two-tailed independent t-tests, 300 lux DI vs FD DI, p = 0.070; Fig. 4F), further providing evidence of an intrinsic diurnal circadian pattern of locomotor activity in the absence of external timing cues. Collectively, these results suggest that NGRs maintain a significant diurnal rhythm across various lighting conditions, and that exposure to bright light increases daytime activity as well as measures of diurnality.

### 3.3 Sex Differences and Individual Variation in Response to Altered Lighting

We next tested for sex differences in behavioral response to light alterations. Comparison of hourly differences and daily averages between 300 and 1000 lux activity reveals differences in how males and females modify their activity profiles when exposed to bright light. In comparing hourly averages from the daily activity profiles, males significantly increase daytime activity in 1000 lux light for 6 of the 12 daylight hours (two-tailed independent t-tests, p<0.05 for ZT 1-5, 7; Fig. 5A), whereas females significantly increase daytime activity in 1000 lux light compared to 300 lux for only 3 of the 12 daylight hours (two-tailed independent t-tests, p<0.05 for ZT 5, 7, 10; Fig. 5A). Plotting daily averages of lights ON and OFF activity across the months-long experiment also reveals that by eye males show sustained increases in activity in the 1000-ON phase, whereas the females show only small increases in activity in 1000-ON days and small decreases in activity during 1000-OFF portion of bright days (Fig. 5B). To analyze this, we compared group daily means (mean activity across all animals of each sex for each day of recording) and show that males had significantly higher 1000 lux daytime activity than females (640,040 ± 61,094 vs 389,292 ± 71,679 pixels/day; t(57) = 14.48, p < 0.001; Fig. 5C). To further compare daily activity levels across all days of the experiment between lighting conditions, we used a Linear Mixed Effects (LME) model (see methods for details). Both males and females significantly increased their daytime 1000 lux activity levels compared to 300 lux, (males: β = 327,875 pixels, t = 13.50, p < 0.001; females: β=63,482, t=4.3, p<0.001; Fig. 5D). No significant effect of lighting condition was observed for male nighttime activity (β = −7,770, t = −0.74, p = 0.46; Fig. 5D), whereas females show significantly suppressed nighttime activity (β=-39,044, t=-4.4, p<0.001; Fig. 5D).

**Figure 5:**
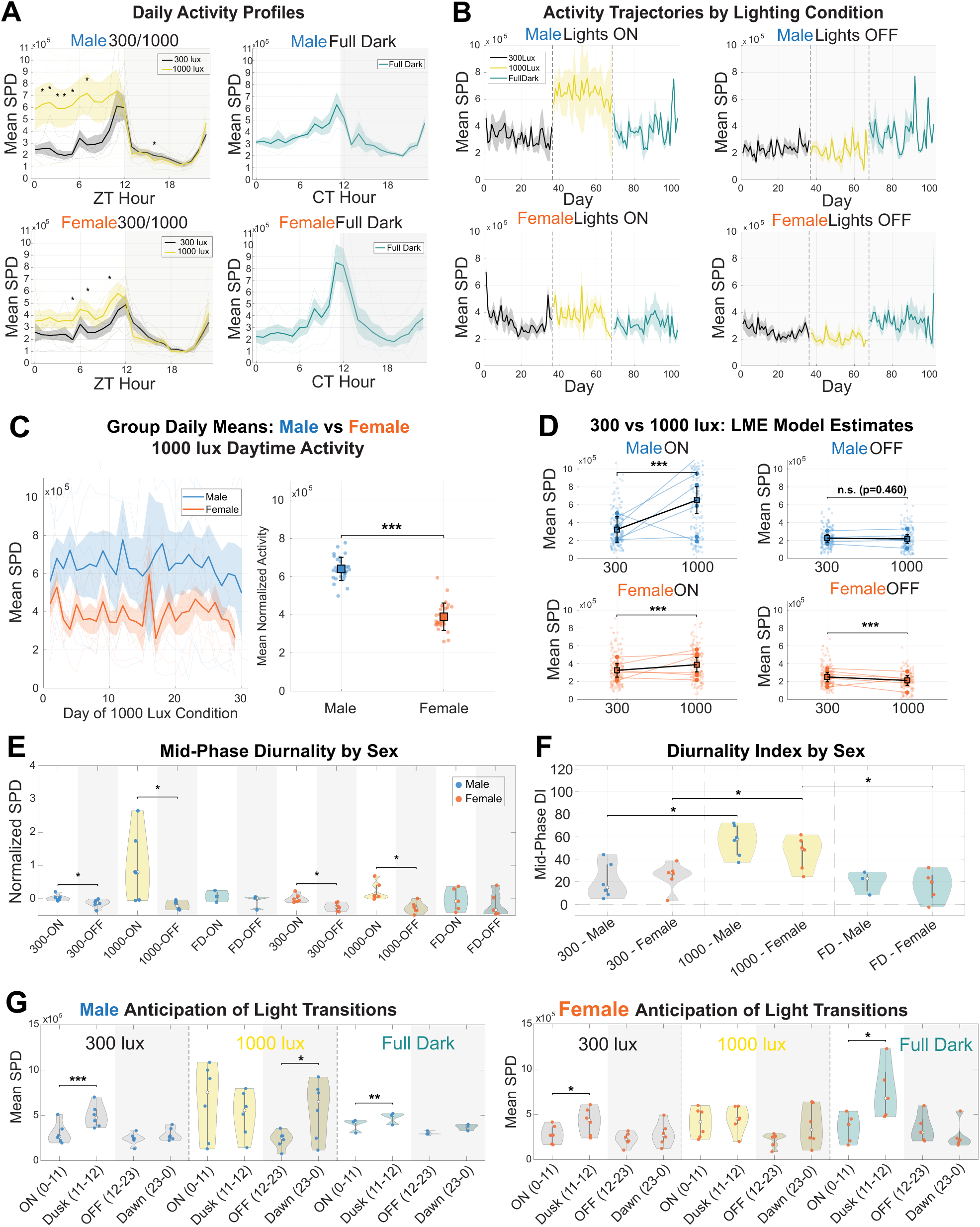
Sex differences in behavioral response to altered lighting conditions. A) Left: Line plots of hourly averages of sex-specific SPD for the 300 lux and Bright conditions. Male (top) and female (bottom) data plotted separately. Asterisks indicate significant differences between hourly means between the 300 and 1000 lux condition. Males appear to show a larger peak at hour 12 under 300 lux than females, and more notably, appear to show a greater increase in Bright-ON conditions than females. Right: Full dark activity profile for all animals (n=8) in the last 7 days of the full dark condition. Each individual animal’s hourly means were re-aligned to CT time prior to averaging across animals by sex. Pooled hourly means are represented by thick line with SEM shading. B) Full lights-ON and lights-OFF daily averages of SPD by sex across all days of each lighting condition. Full dark lights ON and OFF use CT times to better represent subjective day/night. C) Left: Daily group mean activity (± SEM across animals) during the 1000 Lux condition for male (blue) and female (orange) animals across all recording days. Each bold line represents the group mean computed by averaging across all 6 animals of that sex on each day, with shading indicating ± SEM. Faint individual lines show per-animal daily means. Males showed significantly higher daytime activity than females across all 30 days of recording. Right: Distribution of group daily means for male and female animals. Each point represents one day’s group mean (N = 30 male days, N = 29 female days). Squares indicate the mean ± SD across all days. Males showed 64% higher mean daytime activity than females. D) Linear mixed effects model estimates of daytime and nighttime activity at 300 Lux versus 1000 Lux by sex. Four panels show activity comparisons between 300 Lux and 1000 Lux for male daytime, male nighttime, female daytime, and female nighttime. In each panel, faint dots show individual animal-day observations, and lines connect per-animal condition means showing individual trajectories. Large squares with error bars show the LME model-estimated condition means ± 95% confidence interval, with the two model estimates connected by a bold line. The LME model included lighting condition as a fixed effect and individual animal as a random intercept to account for between-animal variability in baseline activity. Significance of the condition effect: *** p < 0.001, n.s. not significant. E) Quantification of mid-ON and mid-OFF mean SPD by light condition and sex (ZT 3-9 for lights ON, ZT 15-21 for lights OFF; full dark CT 3-9 for subjective day and CT 15-21 for subjective night) to emphasize diurnality across conditions. Male (blue) and female (orange) data is pooled per violin, and statistically significant differences between groups are represented by asterisks. Note that male 1000-ON activity is not significantly different than female 1000-ON, likely due to the high variability in the male response. Overall, statistical significances mirror those found in sex-pooled findings in prior figure. F) Diurnality Index across lighting conditions and split by sex (male = blue; female = orange). Note that DI significantly increases in bright light for both sexes. G) Quantification of anticipatory peaks of activity during light-transition times (last hour before transition and first hour after transition, ZT/CT 11-12 and ZT/CT 23-0) compared to averages of 12-hour lights ON or OFF times (ZT/CT 0-11 and ZT/CT (12–23) split into two plots by sex. Note that only males show significant increases in activity in Bright-Dawn compared Bright-OFF, demonstrating that males ramp up activity more than females prior to transitioning back into the light phase.

We then examined degree of increase from 300 to 1000 Lux. Though the effect of 1000 lux light during the daytime was substantially larger in males than females (100% increase in males, 17% increase in females; Fig. 5D), the LME test for sex differences failed to reach significance (β = 260,864, t = 1.72, p = 0.087), likely due to the large variance, particularly in the males (between-animal SD = 271,697 pixels; Fig. 5D). This is likely due to the variability of individual responses in the males, with 4 of the 6 males showing increases in activity in 1000-ON (independent t-tests, male 1000-ON vs male 300-ON, p<0.001 for animal AO1, AO3, AO7, AO9; Fig. S3, Table 3), while 2 of the males surprisingly show no change or a significant decrease in activity (Fig. S3, Table 3). Females show more modest changes in activity in response to the 1000-ON condition, with 2 females showing significantly increased activity, 1 significantly decreased activity, and 3 showing no change (Fig. S3, Table 3).

**Table 3:** 300-ON vs 1000-ON Activity by Sex.

| Animal ID | Sex | N_300 | 300 lux - Mean SPD | N_1000 | 1000 lux – Mean SPD | Delta | t-stat | p-value | Significance |
| --- | --- | --- | --- | --- | --- | --- | --- | --- | --- |
| AO1 | M | 35 | 221,784.4 | 29 | 587,536.1 | 365,751.7 | -9.77 | <0.001 | ***Increase |
| AO2 | M | 35 | 210,540.0 | 31 | 240,605.9 | 30,065.9 | -1.37 | 0.175 | NS |
| AO3 | M | 35 | 192,106.4 | 31 | 821,531.2 | 629,424.8 | -18.75 | <0.001 | ***Increase |
| AO4 | F | 35 | 323,175.2 | 32 | 275,867.2 | -47,308.0 | 2.134 | 0.037 | *Decrease |
| AO5 | F | 36 | 393,240.5 | 26 | 560,229.4 | 166,988.9 | -5.482 | <0.001 | ***Increase |
| AO6 | F | 36 | 305,080.2 | 26 | 298,332.0 | -6748.2 | 0.264 | 0.792 | NS |
| AO7 | M | 36 | 458,196.3 | 26 | 950,209.3 | 492,013.0 | -11.67 | <0.001 | ***Increase |
| AO8 | F | 36 | 476,634.0 | 26 | 507,894.8 | 31,260.8 | -0.831 | 0.410 | NS |
| AO9 | M | 31 | 364,020.7 | 29 | 1,123,896.9 | 759,876.2 | -17.341 | <0.001 | ***Increase |
| AO10 | F | 31 | 212,000.5 | 29 | 488,949.9 | 276,949.4 | -7.142 | <0.001 | ***Increase |
| AO11 | F | 31 | 244,764.4 | 29 | 219,141.7 | -25,622.7 | 0.639 | 0.525 | NS |
| AO12 | M | 31 | 504,854.2 | 29 | 191,988.7 | -312,865.5 | 6.661 | <0.001 | ***Decrease |

Animals in full dark display individual variation in daily activity, as shown in Figure 4, so we next tested if there were activity differences by sex. The daily activity profiles are remarkably similar in structure to the activity profiles seen in 300 and 1000 lux conditions, with two main peaks of activity near expected light transition times even in the absence of light (CT 11 and 23; Fig. 5A). This may be due to the fact that even without light entrainment, male and female free running periods remain close to 24 hours (male tau, mean = 24.07, std=0.93, N=3; female tau, mean=23.80, std=0.35, N=5), and were not significantly different during the final 7 days in full dark (independent t-test, male vs female FD tau, p = 0.562; Fig. S4). Further, group daily means testing showed that neither sex had differences across all days of FD-ON daily activity compared to the 300-ON condition (independent t-tests, 300-ON vs. FD-ON, male: t(61) = -1.803, p=0.0764, female: t(69) = 1.212, p = 0.230; Fig. 5B), representing a return to baseline activity after coming out of the 1000 lux condition.

In Figure 4, we showed that diurnality is maintained in various lighting conditions, and that bright light improves diurnality, prior to differentiating by sex. We next tested whether both male and female NGRs show clear diurnal patterns of activity across lighting conditions. Due to anticipatory activity during light transitions, as shown in Fig. 4A/G, we excluded light transition times and compared average mid-day (ZT 3-9) and mid-night activity (ZT 15-21). Quantification reveals significantly increased activity during the day compared to night for both sexes in both 300 and 1000 lux conditions (paired t-tests; male 300 mid-ON vs mid-OFF, p = 0.02; male mid-1000-ON vs mid-OFF, p = 0.01; female 300 mid-ON vs mid-OFF, p = 0.002; female mid-1000-ON vs mid-OFF, p = 0.007; Fig. 5E). Interestingly, tests for male and female diurnality in the full dark condition during the final 14 days of each condition showed that neither sex alone had significantly more subjective daytime activity than subjective night activity (paired t-tests, male mid-day vs mid-night, p = 0.115; female mid-day vs mid-night, p = 0.080; Fig. 5E), though this is likely due to the decrease in testing power when separating by sex (n=3 male, n=5 female), as the pooled sex data showed that animals in full dark still follow a diurnal rhythm during the fourth week of full dark (Fig. 4E). In comparing the sexes during full dark, we found that males and females show no difference in subjective daytime activity (independent t-test, male vs female FD subjective day, p = 0.813; Fig. 5E) or subjective night activity (independent t-test, male vs female FD subjective night, p = 0.966; Fig. 5E), indicating that both sexes show similar activity levels in FD.

Though the maintenance of diurnality is not sex-specific, we next tested whether exposure to altered lighting differentially affects degree of diurnality (DI) in males and females. Both male and female NGRs show significantly improved diurnality scores in the bright light conditions (paired t-tests, female mid-phase DI 300 vs female mid-phase DI 1000, p = 0.041; male mid-phase DI 300 vs male mid-phase DI 1000, p = 0.020; Fig. 5F), but fail to show sex-specific significant differences (independent t-tests, male vs female mid-phase DI, p > 0.2 for 300 and 1000 lux; Fig. 5F). This suggests that both NGR males and females exhibit strong patterns of diurnal activity irrespective of changes to intensity of environmental lighting, and that bright lighting increases diurnality, though they achieve this in different ways: males show a robust increase in daytime activity in 1000 lux lighting without modulating night activity, while females show more modest increases in 1000 lux daytime light paired with modest decreases in night activity.

In full dark, neither males nor females show significant differences in diurnality scores compared to 300 lux (paired t-tests, male 300 lux DI vs FD DI: p=0.177 (n=3); female 300 lux DI vs FD DI: p=0.459 (n=5); Fig. 5F), and there were no sex-specific differences in diurnality index (independent t-test, FD-DI male (n=3) vs female (n=5), p = 0.718; Fig. 5F). These results suggest that males and females confined to full dark maintain a framework of daily activity with a consistent daily running period and anticipatory day/night transition activity that matches their activity level and patterns in 300 lux.

We then investigated whether activity increased prior to the transition into daytime lights-on (dawn) or nighttime lights-off (dusk) by quantifying activity in the hour before each transition. Both males and females show significant increases in activity at dusk when in 300 lux and full dark conditions (paired t-tests, male 300-ON vs dusk: p=0.0005, male FD-ON vs dusk: p=0.004, female 300-ON vs dusk: p=0.045, female FD-ON vs dusk: p=0.013; Fig. 5G), but this anticipatory activity prior to the lights-OFF transition is not different than daytime activity for either sex in 1000 lux. This homogeneity of activity across ON-hours in 1000 lux is not because the dusk activity drops (paired t-tests, male 300-dusk vs male 1000-dusk: p=0.783, female 300-dusk vs 1000-dusk: p =0.303; Fig. 5G), but rather an effect of increased daytime 1000 lux activity that matches the high activity normally only seen during the dusk transition (paired t-tests, male 1000-ON vs dusk: p=0.137, female 1000-ON vs dusk: p=0.462; Fig. 5G). Interestingly, in the 300 lux and full dark conditions, neither males nor females show significant increases near dawn (paired t-tests, male 300-OFF vs 300-dawn: p=0.129, male FD-OFF vs FD-dawn: p=0.053, female 300-OFF vs 300-dawn: p=0.163, female FD-OFF vs FD-dawn: p=0.094; Fig. 5G), indicating that these animals show less anticipatory activity for the transition into the lights-ON phase; however, males significantly increase dawn activity compared to lights-OFF during the last 14 days of the 1000 lux condition (paired t-tests, male 1000-OFF vs dawn, p = 0.022; Fig. 5G), while females trend towards increasing dawn activity (female 1000-OFF vs dawn, p=0.0645; Fig. 5G) with no significant differences between sexes (male vs female 1000-dawn, p=0.382; Fig. 5G). These results indicate that in standard vivarium lighting and full dark conditions, both sexes increase activity during the end of the lights-ON period in anticipation of lights-OFF, but that exposure to the 1000 lux lighting ameliorates this late daytime burst of activity. Further, in the standard 300 lux and full dark conditions, both sexes fail to ramp-up activity during the end of the lights-OFF phase in anticipation of lights-ON, indicating that at the end of nighttime, only animals exposed to bright 1000 lux lighting during previous daytime begin to raise activity levels in anticipation of daytime.

## 4 Discussion

In this study we use automated monitoring systems to show that in a diurnal rodent, brighter light induces increased daytime-specificity of activity both sexes, but in varying patterns by sex.

The automated monitoring system involves pixel-based actigraphy tracking for single-housed rodents. This tracking system is efficient, able to be automatically processed in batches, and can be seamlessly applied across animals within the same vivarium conditions. Pixel-based actigraphy offers the advantage of continuous recording, higher spatial resolution, more dense sampling, and is more sensitive to pixel gradient changes compared to other tracking methods (i.e. infrared sensors). Many video-based software programs offer similar advantages but are expensive and can be overly complex for basic tracking needs, whereas our tracking system is simple and freely available as open-source Python code. The present study demonstrates the effectiveness of using this system for characterizing in-cage locomotor activity in rodents.

Many rodent models are nocturnal, a trait shaped by evolutionary pressures to avoid predators and exploit nighttime niches. Diurnal rodents like the NGR have evolved adaptations (e.g., visual systems, hormonal rhythms, behavior) for daytime activity. This study helps clarify which aspects of light’s effect on brain and behavior are universal versus those that diverge in diurnal vs. nocturnal species.

Evidence that these animals all show a near 24-hr daily rhythm after a month in full dark conditions indicates a strong endogenous (genetically encoded) circadian rhythm—an evolutionary adaptation favoring activity during daylight. The anticipatory light transition peaks that persist in full dark also promote the idea that these animals are naturally primed to increase activity when transitioning into light. Though our creation of circadian time by manual re-alignment of daily activity profiles produces artificial summation of peak activity prior to lights OFF, the individual pre-shifted profiles all show near 12-hr separation between distinct peaks of activity, and the secondary peak, which is not forced into alignment, persists after pooling CT profiles across animals. This further suggests that animals inherently maintain daily rhythms of activity in expectation of light changes.

Our data provides evidence that NGRs display features of both diurnality and crepuscularity in standard vivarium conditions; however, these animals show clear diurnality after removing the transition phases from the analyses and comparing mid-phase averages. Further, in 1000 lux light conditions, these animals lose their crepuscular-like anticipatory increase in activity (Fig. 4G and 5G), which may indicate that the anticipatory movement in our data may be an effect of the type of artificial lighting in standard vivarium conditions, resulting in what appears to be crepuscular behavior. It’s possible that standard vivarium lighting may insufficiently promote diurnality, induce a more crepuscular pattern, or could reflect that the low-level lighting is sufficient to induce a depressive state in some of the animals, similar to SAD in humans.

Exposure to bright light improves diurnality scores for both males and females, though the data suggests that the male response in 1000 lux conditions is much more robust and variable compared to females, whereas females both modestly increase daytime activity and modestly decrease nighttime activity. This sex-specific behavioral modulation is further demonstrated by hourly activity profiles in the 1000 lux condition which show marked increases in male activity, especially during the first 6hrs of daytime light. The observation that males and females respond differently to lighting could reflect evolutionary divergent reproductive and social strategies as female rodents may be more sensitive to environmental/social context due to demands of reproduction, caregiving, or predator avoidance, while males may be driven by territorial or mate-seeking behaviors. Recent evidence has shown that lesions in the dorsal raphe nucleus selectively increase female daytime activity irrespective of the estrus cycle, which suggests there are sex-specific differences in how the same brain region exerts circadian control of movement (51), and offers insight into why males and females respond differently to the 1000 lux light in our study.

It’s also possible that the high variability in the male response to 1000 lux light, compared to females, is a result of the artificial lighting in an unnatural vivarium setting. In this study we followed the typically used >1000 lux threshold for separating indoor and outdoor lighting, with indoor levels generally measuring in the range of 100-500 lux (52). Our bright light condition may not have been sufficient intensity for all males to alter their activity, considering that outdoor light intensities can range from 1,000 - 100,000+ lux depending on time of day, cloud cover, and environmental surroundings (53–55), though the experienced peak intensities generally measure less than 15,000 lux for the brightest days due to lens filtering properties, and average closer to 1000-5000 lux depending on other environmental conditions (55). Further, most commercially available LED lights are restricted in light emission to certain wavelengths, with many of the missing wavelengths having high relevance for grass rats (43), unlike natural sunlight that encompasses a broad spectrum of wavelengths. It is also unknown if male and female NGRs are differentially sensitive to light, thus, future studies using higher intensity lighting or covering a broader spectrum of wavelengths may better discriminate sex differences in response to bright light.

There is also a possibility that sex-specific responses to 1000 lux light exposure (male individual variability, female attenuation compared to male, and robustness of male activity) are due to stress, lack of social interaction, or depression, which themselves may differentially affect males and females. For example, female daily activity averages over the months-long experiment indicates that females generally decrease activity in the lights-ON phase over time, both from day 0 versus 90 and first vs last day of each condition (Fig. 5B). A new design with lighting condition order reversed would be needed to test this, along with measures for stress hormones and other behavioral indicators of depression. Further, the highly variable male response to the 1000 lux lighting could also indicate individual behavioral phenotypes (i.e. responders and non-responders) often seen with therapeutic intervention for depression, such as antidepressants and light therapy. Whereas the robust activity increase under 1000 lux light in most of the males, but not females, could indicate an evolutionarily preserved mechanism for satisfying sex-specific survival pressures.

Taken collectively, this work emphasizes the importance of further investigating potential sex differences in response to light especially in models sharing diurnality with humans to better understand and treat circadian-associated neuropsychiatric disorders.

## 12 Data Availability Statement

The data analyzed in this study are available from the corresponding author upon request.

## 16 Author Contributions

JH: Writing – original draft, Writing – review and editing, Data curation, Formal analysis, Investigation, Methodology, Project administration, Software, Supervision, Visualization; NM: Writing – review and editing, Data curation, Software, Formal analysis, Visualization; MKM: Writing – review and editing, Software; KLD: Writing - review and editing, Resources; DZ: Writing - review and editing; NO: Writing - review and editing; LY: Writing - review and editing, Conceptualization, Funding acquisition, Methodology, Project administration, Resources, Supervision; BW: Writing - review and editing, Conceptualization, Funding acquisition, Methodology, Project administration, Resources, Supervision.

## Supplementary Material

**Supplementary Figure 1.**
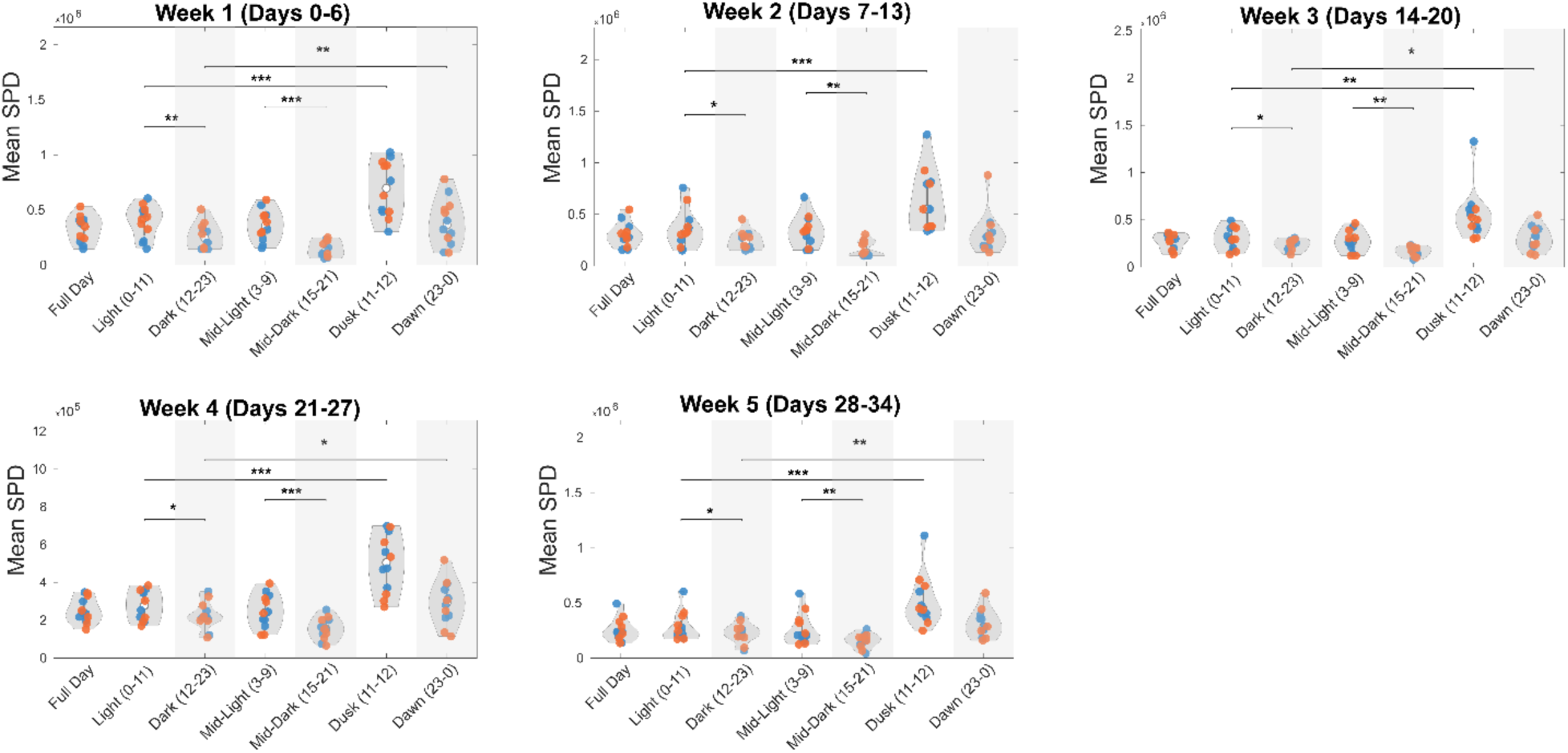
Weekly diurnality testing in standard vivarium lighting. Quantification of weekly mean normalized SPD for comparison of different portions of the day (blue circles = male animals; orange = female). Each plot represents a different week in the 300 lux baseline condition. Weekly separation demonstrates stability of diurnality in the baseline condition.

**Supplementary Figure 2.**
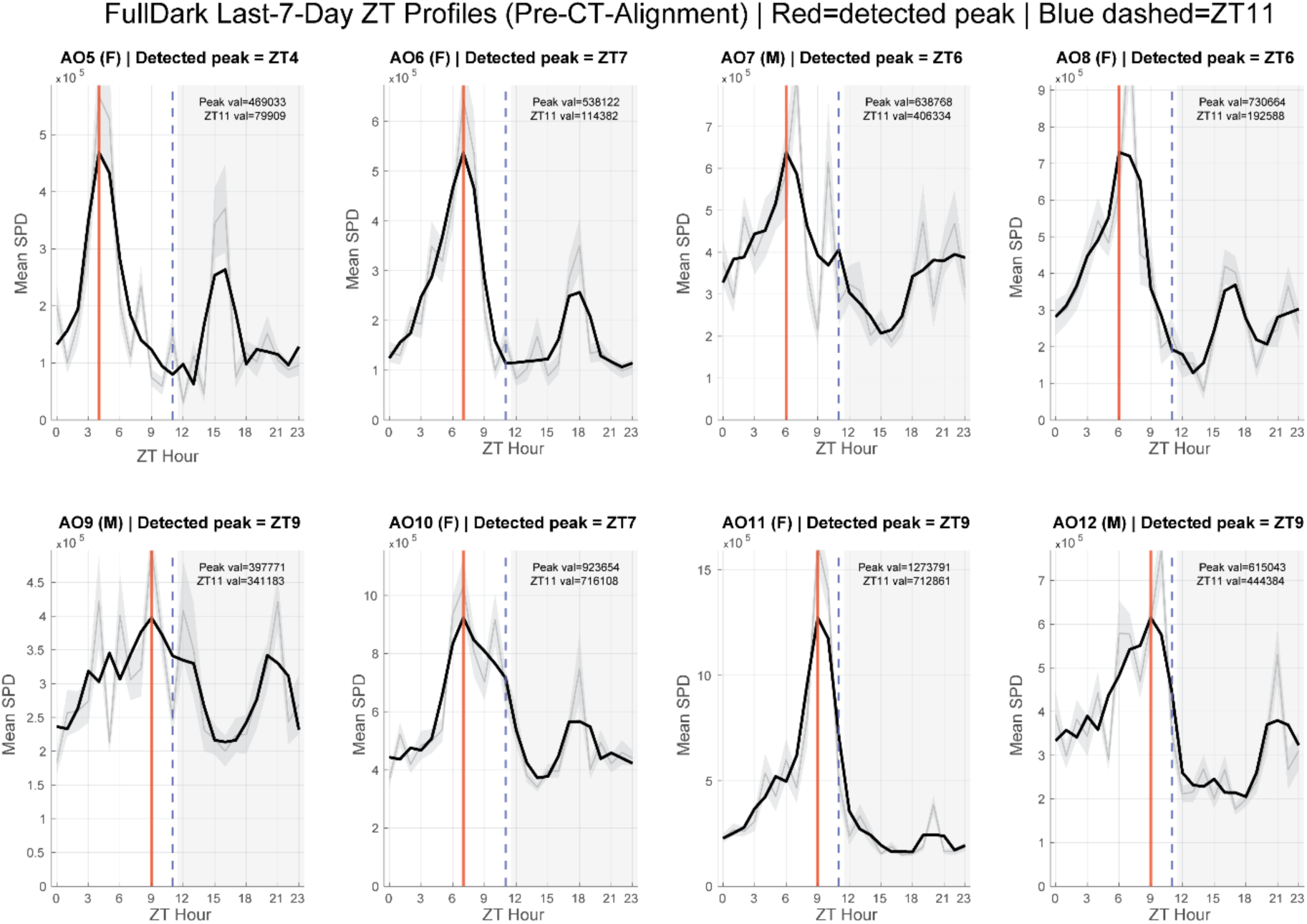
Daily activity profiles prior to re-alignment for Circadian Time. Creation of a new circadian time (CT) was performed by first plotting daily activity profiles for each animal composed of hourly averages across ZT time for the last 7 days in full dark. The black line shows the average SPD by hour and smoothed using a 3-hr moving average. The grey line represents the raw hourly average. The red vertical line indicates the peak activity of each animal and serves as the anchoring point for re-alignment of each animal’s hourly data by defining the peak as the new CT 11 (blue dashed line) to match the ZT 11 peak seen in both male and female data in light-entrained conditions (Fig. 3-5).

**Supplementary Figure 3.**
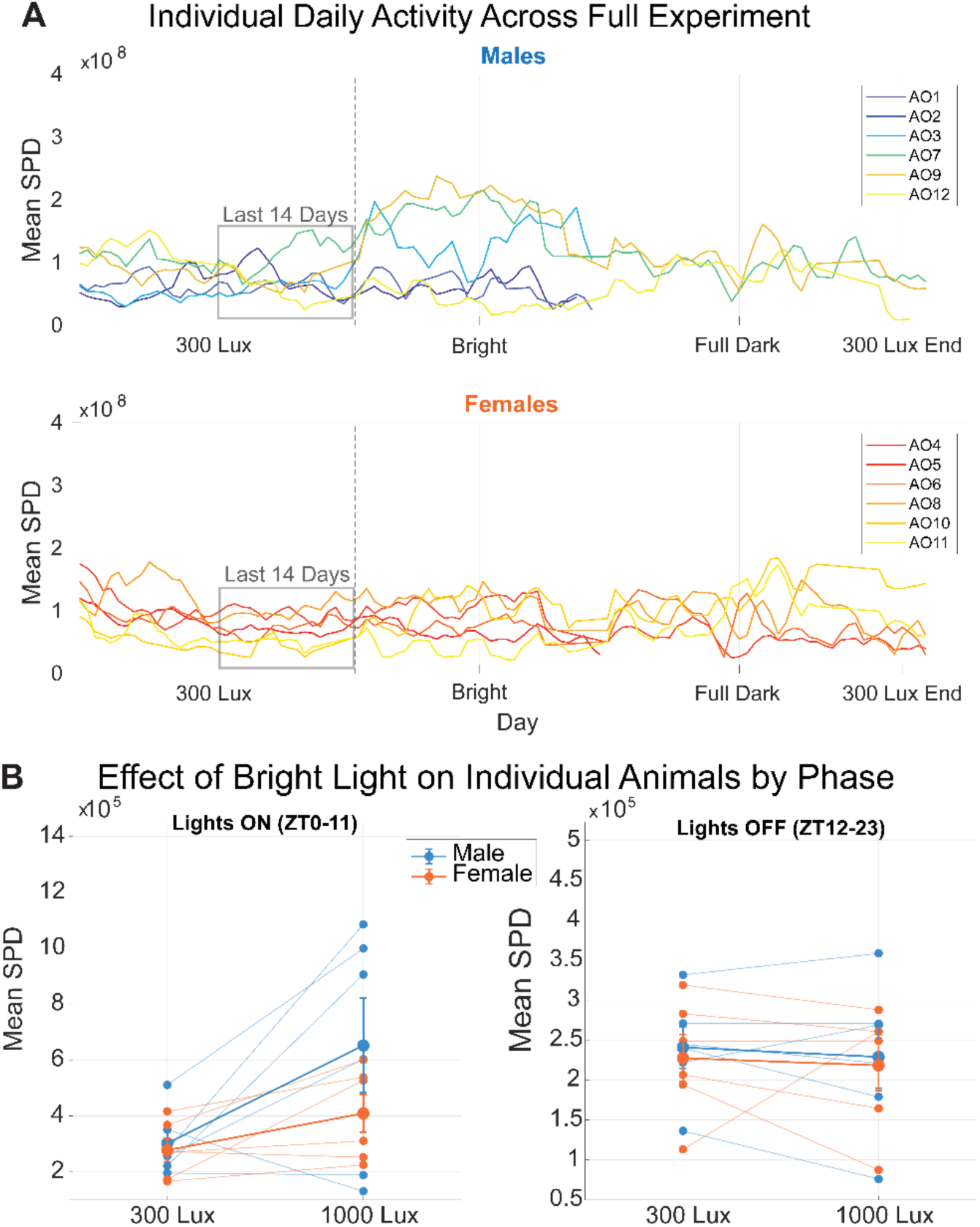
Individual variability in male activity following exposure to bright lights. (A) Individual animal daily mean SPD across months-long experiment (top: males, bottom: females). Data is smoothed using a 3-day moving average. Last 14 days was chosen as the analysis time window based on settling of animal movement after the first 2 weeks, particularly in females (grey box). Notice the activity increase in the males in response to the Bright condition as well as the individual variability in the male response compared to female. (B) Quantification of daily averages from the last 14 days of 300 lux and Bright conditions to show the individual trajectories of each male and female in response to the Bright condition. Notice that in lights-OFF both male and female variability is similar, but in lights-ON males show both larger increases and larger decreases in trajectories than the females.

**Supplementary Figure 4.**
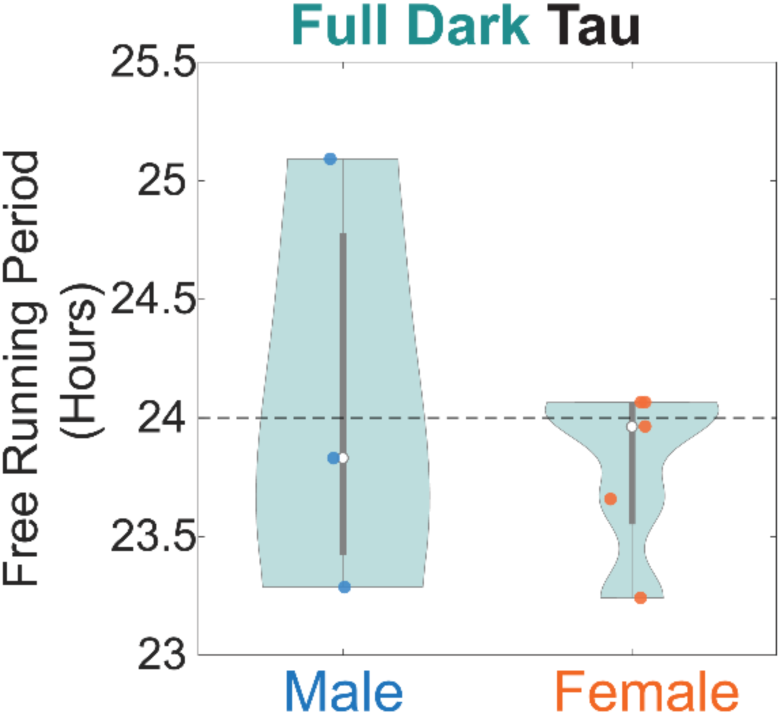
Free running period (tau) by sex. Free running period (tau) computed using a Lomb-Scargle Fourier transform during the final 7 days of the full dark condition by sex (n=3 male, n=5 female). Dashed line indicates the expected 24-hr rhythm. No significant differences are found either between sexes or from the 24-hr daily rhythm; however, notice that some animals experience subjectively longer days and some shorter, forming the basis for the need to re-align daily activity profiles for each animal individually using CT time.

## References

1. Fujisawa S, Amarasingham A, Harrison MT, Buzsáki G. Behavior-dependent short-term assembly dynamics in the medial prefrontal cortex. Nat Neurosci. 2008;11(7):823–33.

2. Manning EE, Dombrovski AY, Torregrossa MM, Ahmari SE. Impaired instrumental reversal learning is associated with increased medial prefrontal cortex activity in Sapap3 knockout mouse model of compulsive behavior. Neuropsychopharmacology. 2019;44(8):1494–504.

3. Ferenczi EA, Zalocusky KA, Liston C, Grosenick L, Warden MR, Amatya D, et al. Prefrontal cortical regulation of brainwide circuit dynamics and reward-related behavior. Science. 2016;351(6268):aac9698.

4. Kim H, Ährlund-Richter S, Wang X, Deisseroth K, Carlén M. Prefrontal parvalbumin neurons in control of attention. Cell. 2016;164(1):208–18.

5. Vogel M, Braungardt T, Meyer W, Schneider W. The effects of shift work on physical and mental health. J Neural Transm. 2012;119:1121–32.

6. Costa G. Shift work and health: current problems and preventive actions. Saf Health Work. 2010;1(2):112–23.

7. Swanson LM, Burgess HJ, Zollars J, Todd Arnedt J. An open-label pilot study of a home wearable light therapy device for postpartum depression. Arch Womens Ment Health. 2018;21:583–6.

8. Burgess HJ, Fogg LF, Young MA, Eastman CI. Bright light therapy for winter depression—is phase advancing beneficial? Chronobiol Int. 2004;21(4–5):759–75.

9. Monteleone P, Martiadis V, Maj M. Circadian rhythms and treatment implications in depression. Prog Neuropsychopharmacol Biol Psychiatry. 2011;35(7):1569–74.

10. Wirz-Justice A. Diurnal variation of depressive symptoms. Dialogues Clin Neurosci. 2008;10(3):337–43.

11. Rosenthal NE, Sack DA, Gillin JC, Lewy AJ, Goodwin FK, Davenport Y, et al. Seasonal affective disorder: a description of the syndrome and preliminary findings with light therapy. Arch Gen Psychiatry. 1984;41(1):72–80.

12. Terman M, Terman JS. Light therapy for seasonal and nonseasonal depression: efficacy, protocol, safety, and side effects. CNS Spectr. 2005;10(8):647–63.

13. Ognjanovski N, Kim DS, Charlett-Green E, Goldiez E, van Koppen S, Aton SJ, et al. Daily rhythms drive dynamism in sleep, oscillations and interneuron firing, while excitatory firing remains stable across 24 h. Eur J Neurosci. 2025;61(1):e16619.

14. Borbély AA. Sleep and motor activity of the rat during ultra-short light-dark cycles. Brain Res. 1976;114(2):305–17.

15. Shuboni D, Cramm S, Yan L, Nunez A, Smale L. Acute behavioral responses to light and darkness in nocturnal Mus musculus and diurnal Arvicanthis niloticus. J Biol Rhythms. 2012;27(4):299–307.

16. Challet E. Minireview: entrainment of the suprachiasmatic clockwork in diurnal and nocturnal mammals. Endocrinology. 2007;148(12):5648–55.

17. Yan L, Smale L, Nunez AA. Circadian and photic modulation of daily rhythms in diurnal mammals. Eur J Neurosci. 2020;51(1):551–66.

18. Smale L, Lee T, Nunez AA. Mammalian diurnality: some facts and gaps. J Biol Rhythms. 2003;18(5):356–66.

19. Esposito E, Li W, T. Mandeville E, Park JH, Şencan I, Guo S, et al. Potential circadian effects on translational failure for neuroprotection. Nature. 2020;582(7812):395–8.

20. Cederroth CR, Albrecht U, Bass J, Brown SA, Dyhrfjeld-Johnsen J, Gachon F, et al. Medicine in the fourth dimension. Cell Metab. 2019;30(2):238–50.

21. Li S, Zhang X, Cai Y, Zheng L, Pang H, Lou L. Sex difference in incidence of major depressive disorder: an analysis from the Global Burden of Disease Study 2019. Ann Gen Psychiatry. 2023;22(1):53.

22. Altemus M, Sarvaiya N, Epperson CN. Sex differences in anxiety and depression clinical perspectives. Front Neuroendocrinol. 2014;35(3):320–30.

23. Eid RS, Gobinath AR, Galea LA. Sex differences in depression: Insights from clinical and preclinical studies. Prog Neurobiol. 2019;176:86–102.

24. Kim K, Kim J, Jung S, Kim HW, Kim HS, Son E, et al. Global prevalence of seasonal affective disorder by latitude: A systematic review and meta-analysis. J Affect Disord. 2025;119807.

25. Pavlidi P, Kokras N, Dalla C. Sex differences in depression and anxiety. In: Sex Differences in Brain Function and Dysfunction. Springer; 2022. p. 103–32.

26. Silva RC, Pisanu C, Maffioletti E, Menesello V, Bortolomasi M, Gennarelli M, et al. Biological markers of sex-based differences in major depressive disorder and in antidepressant response. Eur Neuropsychopharmacol. 2023;76:89–107.

27. LeGates TA, Kvarta MD, Thompson SM. Sex differences in antidepressant efficacy. Neuropsychopharmacology. 2019;44(1):140–54.

28. Bangasser DA, Cuarenta A. Sex differences in anxiety and depression: circuits and mechanisms. Nat Rev Neurosci. 2021;22(11):674–84.

29. Williams ES, Mazei-Robison M, Robison A. Sex differences in major depressive disorder (MDD) and preclinical animal models for the study of depression. Cold Spring Harb Perspect Biol. 2022;14(3):a039198.

30. Mohammadi S, Seyedmirzaei H, Salehi MA, Jahanshahi A, Zakavi SS, Dehghani Firouzabadi F, et al. Brain-based sex differences in depression: a systematic review of neuroimaging studies. Brain Imaging Behav. 2023;17(5):541–69.

31. Dong D, Pizzagalli DA, Bolton TA, Ironside M, Zhang X, Li C, et al. Sex-specific resting state brain network dynamics in patients with major depressive disorder. Neuropsychopharmacology. 2024;49(5):806–13.

32. Blanchong JA, Smale L. Temporal patterns of activity of the unstriped Nile rat, Arvicanthis niloticus. J Mammal. 2000;81(2):595–9.

33. McElhinny TL, Smale L, Holekamp KE. Patterns of body temperature, activity, and reproductive behavior in a tropical murid rodent, Arvicanthis niloticus. Physiol Behav. 1997;62(1):91–6.

34. Novak CM, Smale L, Nunez AA. Fos expression in the sleep-active cell group of the ventrolateral preoptic area in the diurnal murid rodent, Arvicanthis niloticus. Brain Res. 1999;818(2):375–82.

35. Mahoney MM, Smale L. A daily rhythm in mating behavior in a diurnal murid rodent Arvicanthis niloticus. Horm Behav. 2005;47(1):8–13.

36. McElhinny TL, Sisk CL, Holekamp KE, Smale L. A morning surge in plasma luteinizing hormone coincides with elevated Fos expression in gonadotropin-releasing hormone- immunoreactive neurons in the diurnal rodent, Arvicanthis niloticus. Biol Reprod. 1999;61(4):1115–22.

37. Ashkenazy-Frolinger T, Einat H, Kronfeld-Schor N. Diurnal rodents as an advantageous model for affective disorders: novel data from diurnal degu (Octodon degus). J Neural Transm. 2015;122:35–45.

38. Costello A, Linning-Duffy K, Vandenbrook C, Lonstein JS, Yan L. Daytime light deficiency leads to sex-and brain region-specific neuroinflammatory responses in a diurnal rodent. Cell Mol Neurobiol. 2023;43(3):1369–84.

39. Kessler RC, Petukhova M, Sampson NA, Zaslavsky AM, Wittchen H. Twelve-month and lifetime prevalence and lifetime morbid risk of anxiety and mood disorders in the United States. Int J Methods Psychiatr Res. 2012;21(3):169–84.

40. Sramek JJ, Murphy MF, Cutler NR. Sex differences in the psychopharmacological treatment of depression. Dialogues Clin Neurosci. 2016;18(4):447–57.

41. Kornstein SG, Schatzberg AF, Thase ME, Yonkers KA, McCullough JP, Keitner GI, et al. Gender differences in chronic major and double depression. J Affect Disord. 2000;60(1):1–11.

42. Jalnapurkar I, Allen M, Pigott T. Sex differences in anxiety disorders: A review. J Psychiatry Depress Anxiety. 2018;4(12):3–16.

43. Kim AB, Beaver EM, Collins SG, Kriegsfeld LJ, Lockley SW, Wong KY, et al. S-cone photoreceptors regulate daily rhythms and light-induced arousal/wakefulness in diurnal grass rats (Arvicanthis niloticus). J Biol Rhythms. 2023;38(4):366–78.

44. Leach G, Adidharma W, Yan L. Depression-like responses induced by daytime light deficiency in the diurnal grass rat (Arvicanthis niloticus). PLoS One. 2013;8(2):e57115.

45. Lonstein JS, Linning-Duffy K, Yan L. Low daytime light intensity disrupts male copulatory behavior, and upregulates medial preoptic area steroid hormone and dopamine receptor expression, in a diurnal rodent model of seasonal affective disorder. Front Behav Neurosci. 2019;13:72.

46. Yan L, Lonstein JS, Nunez AA. Light as a modulator of emotion and cognition: Lessons learned from studying a diurnal rodent. Horm Behav. 2019;111:78–86.

47. Soler JE, Robison AJ, Núñez AA, Yan L. Light modulates hippocampal function and spatial learning in a diurnal rodent species: A study using male nile grass rat (Arvicanthis niloticus). Hippocampus. 2018;28(3):189–200.

48. Read SA, Collins MJ, Vincent SJ. Light exposure and physical activity in myopic and emmetropic children. Optom Vis Sci. 2014;91(3):330–41.

49. Ostrin LA. Objectively measured light exposure in emmetropic and myopic adults. Optom Vis Sci. 2017;94(2):229–38.

50. Ulaganathan S, Read SA, Collins MJ, Vincent SJ. Influence of seasons upon personal light exposure and longitudinal axial length changes in young adults. Acta Ophthalmol (Copenh). 2019;97(2):e256–65.

51. Tamogami S, Okeya M, Suzuki R, Amano H, Yamamoto R, Mochizuki T, et al. Sex differences in serotonergic control of daytime activities in diurnal Nile grass rats. Brain Res. 2025;149862.

52. Daugaard S, Markvart J, Bonde JP, Christoffersen J, Garde AH, Hansen ÅM, et al. Light exposure during days with night, outdoor, and indoor work. Ann Work Expo Health. 2019;63(6):651–65.

53. Norton TT, Siegwart Jr JT. Light levels, refractive development, and myopia–a speculative review. Exp Eye Res. 2013;114:48–57.

54. Lou L, Ostrin LA. The outdoor environment affects retinal and choroidal thickness. Ophthalmic Physiol Opt. 2023;43(3):572–83.

55. Lanca C, Teo A, Vivagandan A, Htoon HM, Najjar RP, Spiegel DP, et al. The effects of different outdoor environments, sunglasses and hats on light levels: implications for myopia prevention. Transl Vis Sci Technol. 2019;8(4):7–7.

